# Did circular DNA shape the evolution of mammalian genomes?

**DOI:** 10.1101/2022.06.22.497135

**Authors:** Sylvester Holt, Gerard Arrey Tané, Birgitte Regenberg

**Affiliations:** Department of Biology, University of Copenhagen, Copenhagen DK-2100, Denmark

## Abstract

Extrachromosomal circular DNA of chromosomal origin (eccDNA) can rapidly shape the evolution and adaptation of mitotically dividing cells such as tumor cells. However, whether eccDNA has a permanent impact on genome evolution through the germline is largely unexplored. Here, we propose that a large fraction of the syntenic changes that are found between mammalian species are caused by germline transposition of eccDNA. We have previously shown the existence of eccDNA in mammalian meiotic cells. By reanalysis of available synteny maps, we now find that up to 6% of mammalian genomes might have rearranged via a circular DNA intermediate. Hence, eccDNA in the germline is expected to have large effects on evolution of gene order.

**Highlights:**

- Extrachromosomal circular DNA (eccDNA) is present in mammalian germline cells showing that eccDNAs are not excluded, repressed or eliminated during meiosis
- Large eccDNA reinsertions into the genome can change gene synteny in a recognizable pattern based on its circular junction and integration breakpoint.
- By reanalyzing synteny maps form 8 mammalian species, we show that 6% of genome of the ungulates cow and sheep can be explained by eccDNA insertions.
- We propose that reinsertion of large eccDNA that are fixed in germline cells may have contributed to speciation barriers and evolution of new species

## Introduction

Work over the past decade has demonstrated that eccDNA is common in a wide range of eukaryotic organisms ranging from unicellular yeasts to multicellular plants [1–5] and eecDNA has been found in a wide range of mitotic tissue and cells in sizes large enough to carry whole genes [1,2,6–9] and eccDNA can thereby have a direct effect on cell phenotype if they carry expressed genes and replicate prior to mitosis (see Arrey et al for an overview) [10]. The best described examples from animals are the large extrachromosomal DNA molecules (ecDNA) in tumors with proto-oncogenes, which have been shown to drive tumorigenesis by increasing the copy number heterogeneity of oncogenes and the gene expression levels [8,11–14]. After DNA replication, the lack of centromeres allow the ecDNAs to escape from the symmetric segregation imposed on chromosomes during cell division. As a result, ecDNAs segregate in an uneven fashion, allowing a fraction of cells to accumulate large copy numbers [8,15–17]. Phenotypic effects of circular DNAs are often transient due to their random segregation. However, reinsertion of ecDNA into chromosomes can lead to the formation of arrays of genes, known as homogeneously staining regions (HSR), which fix the selective advantage and create a stable inheritance of the gene amplification in the tumors that carry them [18].

However, whether circular DNA is present in the mammalian germline is largely unexplored despite the many recent studies of other structural variants in germline [19–21]. EccDNA was first described in boar sperm [22] but since then and until recently the presence and impact of eccDNA in germline cells has been overlooked. We and others recently demonstrated the existence of eccDNA from chromosomes in the human germline cells [6,23]. This is interesting because this eccDNA in germline cells could potentially reinsert into the human chromosomes and make stable alterations that are inherited across generations. At a broader level this would imply that transposition of Mb eccDNA provide the basis for changes in gene order (synteny) that is part of genome evolution.

We now propose that major chromosomal rearrangement in the germline can happen through a circular DNA intermediate formed from one part of a genome that reinsert in another part of the genome (the circle translocation theory, Figure 1). For this to happen (i) deletions of chromosomal DNA in germline cells must form circular intermediates, and (ii) occasionally reinsert into chromosomes to form balanced rearrangements with no or little gene loss. Formation and reinsertion must (iii) occur in germline stem cells, cells in meiosis or in zygotic cells shortly after fertilization and subsequently be carried by organisms that go through inbreeding to ensure homozygotes are formed with the new constellation of genes. Several mammals have well annotated genomes with synteny maps that allow this hypothesis to be tested, and their relatively small population sizes can secure rapid fixation of the gene rearrangements in new species. We therefore used mammalian genomes to describe the evidence for eccDNA in germline and explored existing genome maps across the mammalian class to identify regions with synteny patterns that correspond to transposition of circular intermediates. We also review evidence for alternative theories and finally describe how the mammalian genomes and especially the human genome regulates the level of circles translocations.

**Figure 1.**
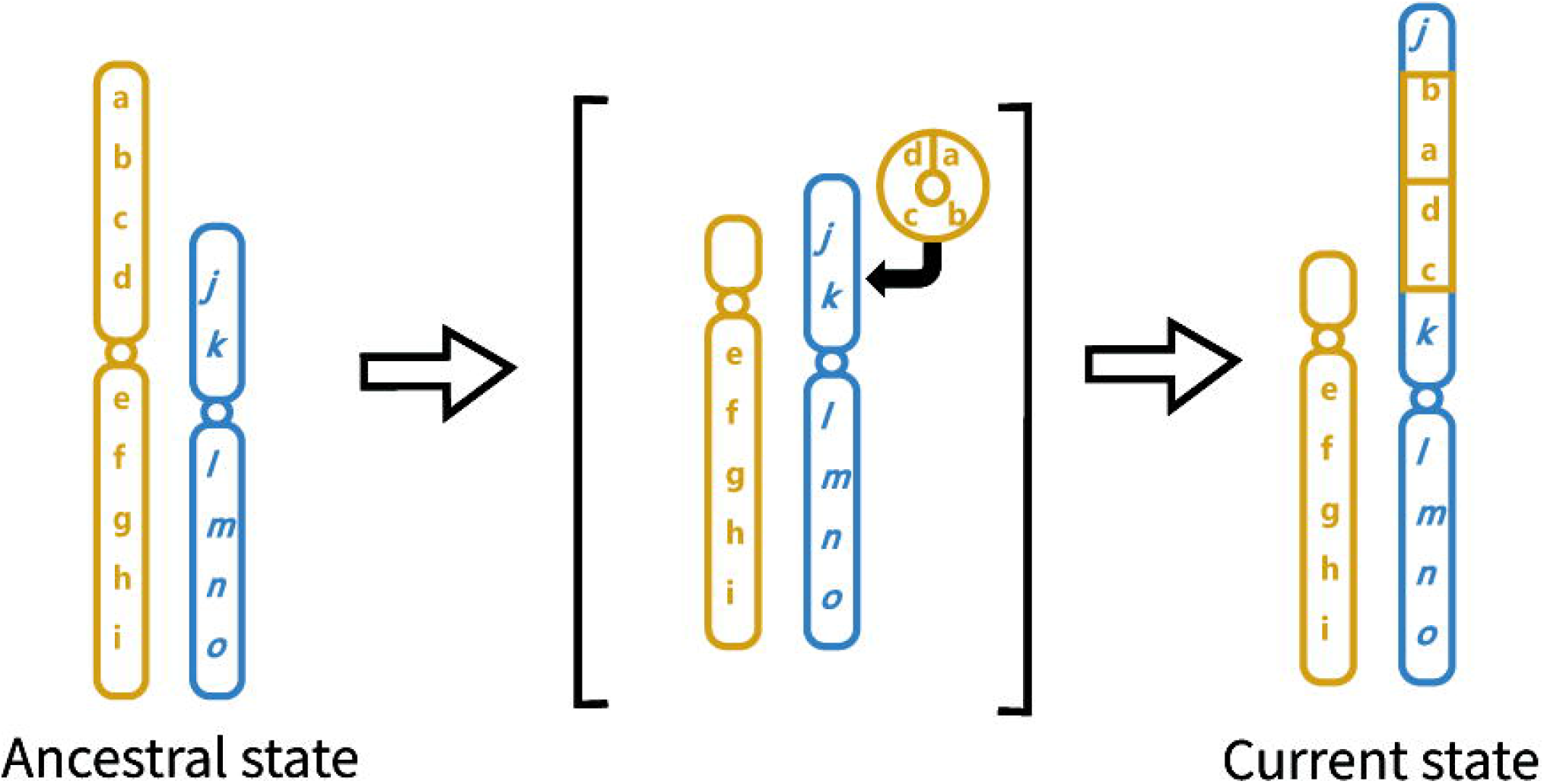
The circle transposition theory. DNA is deleted from a chromosome, circularizes and reinserts in a different position in the genome. The syntenic changes caused by circle re-insertion is designated as a circle insertion pattern. (|abcd| ➡ [abcd^circle^] ➡|ba|dc| or |cd|ab|)

Our results are important because they provide insight into how the mammalian genomes might have evolved through changes in gene order, which might cause reproductive barriers and new species through sterility in inter-species hybrids. Translocation of genes in the genome may also provide new phenotypic features if a gene replacement allows for higher transcription of the genes or expression in new tissues due to new combinations of gene and enhancer elements.

### i) Circular DNA in mammalian germlines

To test if eccDNA is formed during meiosis of mammalian germline cells, we recently isolated and sequenced eccDNA from 1 million human sperm cells of 29 different donors [6]. Using long-read sequencing technology we could detect 8,346 eccDNAs originating from a single chromosomal loci and 3,090 chimeric circles from two or more different chromosomal localizations, thus showing that eccDNA is common in human germline cells. In further support of this, eccDNAs have also been isolated from germline cells in wild boar [22], *Caenorhabditis elegans* [24], in the plant germ cells [4], and more recently in a pooled sample with an unknown number of human donors [23] and in mouse sperm cells [25,26]. In none of these studies were large Mb sized circles found, though this is expected. During meiosis, one double stranded break that recombines with elements on the same chromatid can lead to a deletion and a circle forming through intrachromatidal non-allelic homologous recombination (NAHJ) if the deleted fragment contains sufficient homology. Experimental data from Turner et al. 2008 [27] measured the recombination rates during meiosis in human sperm cells for four loci and found that intrachromatidal non-allelic homologous recombination (NAHJ), was as frequent as recombination chromatids or chromosomes (2*10^−6^ to 4*10^−5^). With such a high potential for circle formation, megabase sized circles are expected to appear. In human cancers, megabase sized ecDNA that contain oncogenes are identified through whole-genome sequencing and FISH microscopy [8,14,28]. The lack of large circles in the studies where eccDNA is isolated from sperm, could therefore be due to technical limitations such as breakage of the large DNA molecules before exonuclease treatment or unspecific cutting with restriction enzymes to remove linear and mitochondrial DNA.

### ii) Reinsertion of eccDNA in germline cells

Evidence of circle reinsertion is mainly found from transposable elements (TE). TE move around the genome aided by a self-encoded specialized machinery depending on their class, which may include different enzymes: retrotranscriptases, integrases and endonucleases as well as helicases and transposases. While doing so, they can change gene expression, gene order and create recombination hotspots in the loci where they integrate [29].

Similarly, but not involving any transposition machinery, eccDNA transpositions might also allow fragments of DNA to translocate and contribute to similar changes. A few example are know from mitotic cells where both circle and the potential insertion product is found. Translocations of genes on circular elements are known from the unicellular eukaryotic yeast cells where circularization, amplification and insertions back into the original locus serve as a mode of environmental adaptation [30–32]. Yeast cells carry two glucose transporter genes, HXT6 and *HXT7*, in direct repeat on chromosome IV that often forms a self-replicating circular intermediate, [HXT6/7^*circle*^]. When exposed to glucose limitation the fraction of cells carrying the [*HXT6/7*^*circle*^] increase for several generation until a stable chromosomal *HXT6 HXT6/7 HXT7* amplification occurs and outcompetes the wild-type *HXT6 HXT7* genotype and the [*HXT6/7*^*circle*^] amplification (Prada-Luengo et al., 2020). This result supports that selective advantages created by the eccDNA intermediate can be fixed in a population through stable insertions. Another example is the artificially introduced XYL locus that form [*XYL*^*circle*^] and stable chromosomal amplifications that might arise from the [*XYL*^*circle*^] amplification [31].

Traces of circle insertion have also been identified in multicellular eukaryotes. Evidence for eccDNA integrations in meiotic cells come from few examples in fish and mammal species. In the Nile tilapia cichlid fish (*Oreochromis niloticus*), amplification of the vasa genes via a circular intermediates has been proposed to explain the genomic re-arrangement of these loci [33]. The original vasa locus was found to be duplicated from the original site and integrated into two different sites, one involving a 28-kb eccDNA intermediate. Because other members of the cichlid family do not share these two insertions, the eccDNA transposition might have occurred during the evolution of the Oreochromis genus [33]. Durkin et al describes how the color sidedness on three cow varieties are associated with syntenic changes that can be associated to circle insertions. The Belgian blue cow has a 492-kb eccDNA insertion in chromosome 29 spanning *KIT* gene from chromosome 6. A portion of the allele further circularized again and a 575-kb fragment of mixed chromosome 6 and 29 inserted back in chromosome 6 in Brown Swiss. These serial translocations affected *KIT* expression leading to different color sidedness patterns in these cow breeds. [34]. Finally, two studies have identified circle insertion patterns between human and a few other primates, but only as a consequence of segmental duplications [35,36].

In humans, eccDNA integrations have been investigated in the context of mitotic cells, in cancer. *EGFR* is an oncogenic growth factor receptor often times amplified within eccDNA molecules in tumors, driving cancer growth [12]. Amplifications of the *EGFR* gene in eccDNA can further be fixed in the chromosomes by integrating copies in arrays also called homogeneous staining regions (HSR) [8]. Another example is the integration of a TRECβ circle into the upstream region of *TAL1* gene, deregulating its transcription levels and causing onset of acute lymphoblastic leukemia [37].

#### Test of the circle transposition theory

To formally test whether circle insertions could have shaped the evolution of mammalian genomes, we analyzed for changes in gene order (synteny) that suggests germline insertions of eccDNA (Figure 2, key figure). Coordinates for synteny blocks were extracted with a 150 kb window using the SynBuilder tool at the Synteny Portal [38] for eight different representative mammalian species. The construction of the synteny maps is described by Lee et al. (2016). Briefly, the whole-genome alignments against the human reference genome (hg38) are merged until the given resolution. From the synteny coordinates, we searched for circle insertion patterns using R code (available in supplementary materials): 1) two flanking inversions, or 2) two flanking blocks in the same direction with a change in order (Figure S1).

**Figure 2.**
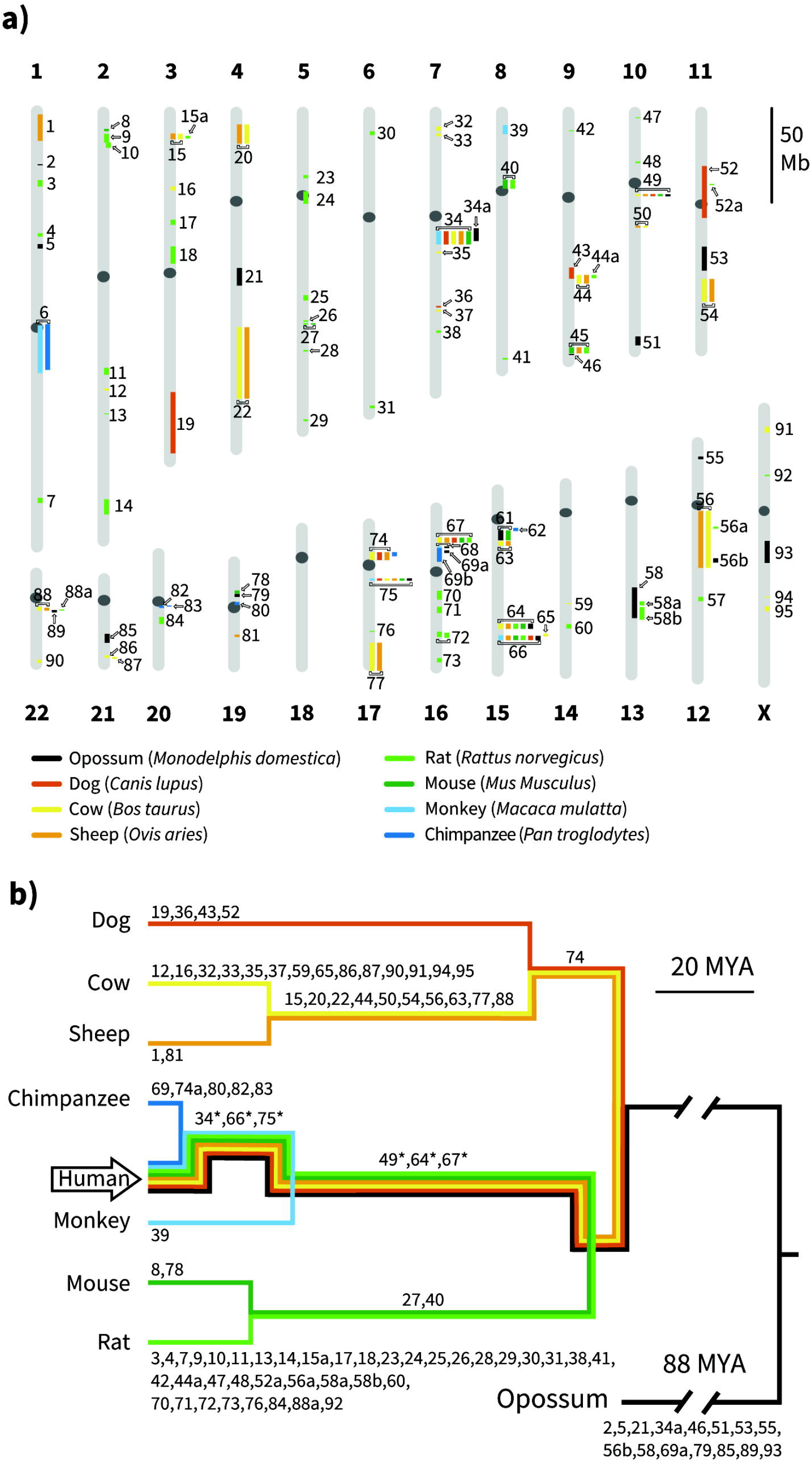
Circle insertion patterns in 8 representative mammals. a) Circle insertion patterns for 8 mammals, projected onto human genome coordinates b) Phylogenetic tree indicating shared and unique circle insertion patterns. The tree is read like a mirror from the human branch (arrow) towards the branches for the remaining organisms. The synteny coordinates between the organisms were extracted with SynBuilder at the Synteny Portal web server [38] and circle insertion patterns were identified with custom made R-script available in the supplementary materials. The tree was generated with TimeTree [52] and colors indicate the organisms that have the patterns, shown in a). The patterns are also indicated with numbers. Star indicate that the pattern may be degenerated in a few organisms.

If the patterns are observed in multiple species, we can infer their position in the phylogeny and the organism in which they are evolved. For example, if the circle insertion pattern is unique for human, we expect all the other species to have the pattern mirrored in the maps, unless the pattern was degraded through other events. We did not observe any human specific events that were found mirrored in all 8 species, which is likely because requiring the synteny to be identical in 8 species is strict, given the evolutionary timescale of ∼159 million years. However, some patterns were still highly conserved in many species, indicating that the event happened in the split of the primates (Figure 2B and Figure S6-S8).

The patterns shown in Figure 2 reveal that major parts of all tested mammalian genome can have evolved through balanced transpositions that are consistent with and circular intermediate (Figure 1). Ungulates, cow and sheep, appeared to have undergone the largest transformation of theirs genomes with 6% being altered compared to their closest associate (Table 1). We confirmed that the synteny patterns were overlapping with orthologous open reading frames for three of the largest patterns in ungulates (Figure S2-S4).

**Table 1.** Characteristics for the synteny maps and circle insertion patterns. The coverage is the total size of the circle insertion patterns divided by the size of the genome. The average size is the average size of the circle insertion patterns identified. Both are listed in coordinates of the respective species.

| Species | Number of blocks in the synteny maps | Number of patterns | Average size of pattern (Mb) | Coverage of genome |
| --- | --- | --- | --- | --- |
| *Monodelphis domestica* | 673 | 21 | 5.0 | 2.9% |
| *Canis lupus* | 343 | 10 | 6.8 | 2.8% |
| *Bos taurus* | 470 | 30 | 5.4 | 6.0% |
| *Ovis aries* | 404 | 20 | 7.8 | 6.0% |
| *Rattus norvegicus* | 882 | 46 | 2.6 | 4.1% |
| *Mus musculus* | 408 | 12 | 2.9 | 1.3% |
| *Macaca mulatta* | 261 | 6 | 3.1 | 0.6% |
| *Pan troglodytes* | 120 | 7 | 2.2 | 0.5% |

### iii) Fixation of the circle insertion

A requisite for genetic mutations to be have an evolutionary impact is to be present in germline cells. Germline mutations can occur either in the meiotic cells right before fertilization or in their gonad precursors during meiotic maturation. They can also occur within the first stages of a zygote after fertilization during zygote development. The number of divisions of the zygote will determine if the mutation is present in all cells or in just a fraction. If the mutation occurs before the somatic and germline split (also named gonosomal mutations) an unbiased fraction of both somatic and germline cells will carry the mutation. If it occurs after the split, only a fraction of either the somatic or germline cells will carry it.

The fact that eccDNAs have been detected in sperm of multiple animals, including boar, mice and human suggests that eccDNAs are not excluded, repressed or eliminated during meiosis. In fact, they have been recently shown to arise via *SPO11* cleavage of chromatids during meiotic recombination in mice, showing that meiosis is a source of eccDNAs in itself [25]. While examples of eccDNA translocation in fish, cows and humans imply a germline transmission [33–36], it is yet not clear when this event occurs during gamete maturation (from the generation of F0 haploid cells to the F1 diploid). While eccDNA insertion within the pre-meiotic (F0, 2n), meiotic, post-meiotic (n) cells or within the zygote (F1, 2n) would certainly lead to a rearranged and homogenous F1 germline cell population, eccDNA integrations occurring within the first zygotic divisions would lead to a mosaic F1 germline cell population.

A second requisite for germline mutations to have an impact in evolution is successive rounds of inbreeding in reproductive isolated populations [39]. From the heterozygote founding event in a single individual, the allele can be fixed in a population (when all members have the allele) due to selective pressure and genetic drift. Because eccDNA translocations are a *priori* balanced (no loss of genetic information), they may result in normal development allowing its phenotypic effects (changes in gene expression) to be expressed. Nonetheless, due to low reproductive fitness of large structural chromosomal heterozygotes, fixation of a novel rearrangement might be rare, reducing their effect in speciation [40]. Some mechanisms, such as inverted meiosis (sister chromatids separate before homolog chromosomes), tolerate such heterozygosity and rescue meiotic fitness [41,42]. Inverted meiosis is more commonly found in species with holocentric chromosomes (kinetochore activity is localized alongside the chromosome) in different phyla of animals and plants [43] but have been also described in monocentric species like humans [44] and yeast [45], suggesting that large chromosomal changes can indeed be tolerated in meiosis of heterozygous hybrids.

#### Alternative theories

Two alternative mechanisms could explain the eccDNA insertion patterns in Figure 2. These are (i) random insertion of two flanking linear DNA fragments in the same location (ii) replication errors (MMBIR). Forming a circle insertion pattern could occur through random insertion of two adjacent linear DNA fragment in sequential events. However, the likelihood of two linear fragments inserting randomly in the same locus flanking each other with inverted direction is presumably low. We tested whether the patterns observed in our study could be due to random insertion two linear fragments by randomizing the synteny blocks for all organisms 10,000 times and extracting circle insertion patterns. In all cases the number of identified patterns in randomized data was much lower than observed in the unscrambled genomes (Figure S10), confirming that random insertion of linear fragment is unlikely to generate circle insertion patterns.

Alternatively, the observed eccDNA insertion patterns could be caused by stalled DNA replication forks that lead to DNA damage and are repaired through strand replacement successive strand replacement in areas with microhomology, termed Microhomology Mediated Break Induced Repair (MMBIR). This theory is used to explain patterns overserved in complex rearrangements in congenital disorders [46,47]. To produce circle insertion patterns similar to the ones we describe in this study (Figure 2), the replicating DNA strand would have to continue replication in another chromosome, then jump upstream in the same chromatid, continue replication until the exact starting position and jump back to the original chromosome and continue synthesis. Because this would require sequence of very precise strand replacements in multiple locations in different chromosomes, we are of the opinion that transposition of an eccDNA involving only two events is a more parsimonious explanation for the patterns seen in Figure 2.

#### Do genomes regulate circle transposition?

While our data suggest that some circle transpositions have been fixed in the genome of mammals, most DNA circularizations and reinsertions might have deleterious effects. Circles forming from gene rich regions are likely to lead to gene deletions and loss of function as most eccDNA are predicted to be lost. Insertion of eccDNA might also have deleterious effect if they insert in genes, enhancer or promoter regions that disrupts an essential function/connection between the coding region of a gene and it enhancers. An example of the consequences are the congenital disorders in humans, that are caused by large complex insertions [27]. It therefore seem likely that the eukaryotic genomes have developed mechanisms to regulate the level of DNA circularization and insertion.

We have previously found a negative correlation between the number of genes and *Alu* on a chromosome and the number of eccDNA formed from the chromosome in human germline cells [6]. Chromosomes with many genes and *Alu* elements, such as chromosome 17 and 19, exhibit the lowest rate of circularization. Why *Alu* elements are conserved on gene rich chromosomes have been an open question since the sequencing for the human genome [48]. In the original paper, it is suggested that there is positive selection for *Alu* elements, and that this is related to gene richness. Our results thereby offer a long anticipated explanation for how *Alu* elements coevolved with genes to protect genome integrity against intra-chromatidal recombination that result in deletions and eccDNA, and thereby support the model that there has been positive selection for *Alu* elements in the human genome.

The *Alu* elements are specific for a small group of mammals, and the correlation between low circularization and high levels of *Alu* elements is therefore not expected to “hold” for other mammals. Still there might be similar cis-acting elements in other mammals that could regulate the level of DNA circularization in gene rich regions of their genomes [49].

Whether reinsertion of extrachromosomal DNA is regulated is less well understood. Lukaszewicz and coworkers recently suggest that circularization of extrachromosomal DNA in the germline may actually reduce the probability that the DNA reinserts into a chromosome [25]. This is backed by early studies of DNA integration of plasmids in yeast where linearized plasmids have much higher rates of insertions than intact circular plasmids [50].

#### Concluding remarks

Mammalian genomes are full of genome rearrangements that can have shape the expression of genes in the elements. However, a key open question has long been how these rearrangements arise at the molecular level. We have proposed a theory that can explain up to 6% of the differences between e.g. ungulates and human. Hence, our model and findings support that transposition of eccDNA have an important role in the evolution of eukaryotes. Our theory can also be applied to test how eccDNA might have shaped the genome of other classes, phyla and kingdoms. eccDNA are also known from plants, both as products of TE and genes [4,51] and it therefore seems plausible that eccDNA may have shaped plant genomes through stable insertions of eccDNA. Because large insertions and deletions can lead to reproductive barriers between individuals with the mutations and the parental genotype, eccDNA transposition might have contributed to reproductive barrier that lead to new species across the tree of life.

## Glossary

Balanced chromosomal rearrangement: A type of chromosomal structural variant (SV) involving chromosomal rearrangements without cytogenetically apparent gain or loss of chromatin.
Extrachromosomal circular DNA (eccDNA): Covalently closed circular DNA without a centromere formed from linear chromosomal DNA.
Large extrachromosomal circular DNA molecules (ecDNA): EccDNA with a size over 1 Mb and visible by light microscopy.
Structural variation (SV): Chromosomal rearrangements including translocations, inversions, and insertions.
Synteny: The physical co-localization of genetic loci on the same chromosome within an individual or species.
Transposition: The transfer of a segment of DNA from one site to another in the genome.
Circle insertion pattern: A term that we introduce here for the first time. Two flanking synteny blocks with inverted direction and order (|abcd| ➡ [abcd^*circle*^] ➡|ba|dc| or |cd|ab|.
Microhomology-mediated break-induced replication (MMBIR): A replication-based mechanism of recombination between sequences with microhomology.

### Outstanding questions

- Did large genome rearrangements lead to reproductive barriers and thereby important for speciation?
- How does circular DNA transposition happen at the molecular level in the window around meiosis?
- Since chimeric (complex) circles are abundant, are there also complex patterns from re-insertion of eccDNA?
- How do gene rich chromosomes in human prevent intra chromatidal recombination that leads to DNA circularization?
- Are eccDNAs that are formed in mitotic cells passed through meiosis allowing them to enter the germline?
- Can environmental stress factors increase the rate eccDNA formation and reinsertion in the germline?
- How particular genomic features (*Alu*, tandem arrays, etc.) modulate circular DNA transpositions?
- Due to reduction of meiotic success of large chromosomal rearrangements, are small eccDNA transpositions more tolerated and more frequent?
- Do eukaryotic phyla tolerate differently eccDNA transpositions?
- To which extend did eccDNA transpositions shape gene architecture (exon/intron)?

