## Supplemental materials for "Did circular DNA shape the evolution of mammalian genomes?"

**Supplementary materials for “Did circular DNA shape the evolution of mammalian genomes?” Holt et al.**

### Extraction of circle insertion pattern in mammalian synteny maps

Coordinates for synteny blocks were extracted with a 150 kb window with the SynBuilder tool at the Synteny Portal [1] for eight different representative mammalian species. The construction of the synteny maps is described by Lee et al. (2016). Briefly, the UCSC genome browser whole-genome alignments against the human reference genome hg38 are merged until the given resolution. From the synteny coordinates, we searched for circle insertion patterns using R code: 1) two flanking inversions, or 2) two flanking blocks in the same direction with a change in order (Figure S1).

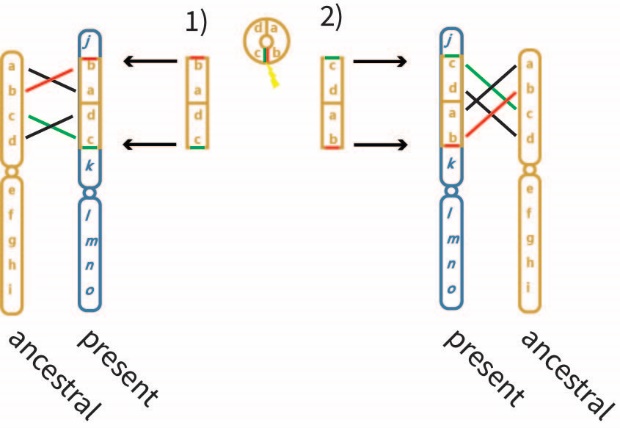

**Figure S1. Circle insertion patterns. The circle breaks open and may integrate in either of two different directions. When comparing the present genome to the ancestral, a circle that inserted may therefore leave a pattern of 1) two flanking inversions, or 2) two flanking blocks with a change in the order.**

### Comparison of patterns found in multiple species

Insertion in either direction (1 or 2) are treated equally in this study. All circle insertion patterns identified are listed in Table S1, with patterns highlighted that are found in multiple species that are less likely to be spurious. In table S2, we have also done a multiple species comparison and the inferred place in the phylogenetic tree based on overlapping patterns. Finally, to check the quality of the patterns for the multiple species, we also provide their annotations (Figure S2-S9). Specific patterns are discussed in the text below.

#### Circle insertion patterns for primates

Here, we show three patterns for primates. Because the patterns is a mirror for all other species, they require all the remaining species to have the pattern at the same position. This means that there is much higher chance that one or more species have degenerated at the exact position over the course of evolution. We therefore consider patterns for primates that have one or two degenerated patterns (designated with * in Figure 1). We discuss three patterns observed for all three primates (64 and 67; Figure S2 and S3) and for human and chimpanzee (66 and 75; Figure S4 and S5).

##### Insertion patterns 64 for primates

The pattern was found opossum, cow, sheep, rat and mouse, but was completely degenerated for dog since the blocks overlapping were linked to three different chromosomes (Figure S2). For cow, the genes in block 2 (*ACSBG1*, *WDR61* , *DNAJA4*, *IDH3A*, *CRABP1*) were found in the flanking block upstream, suggesting that the pattern was conserved, but not correctly identified in the Synteny Portal software.

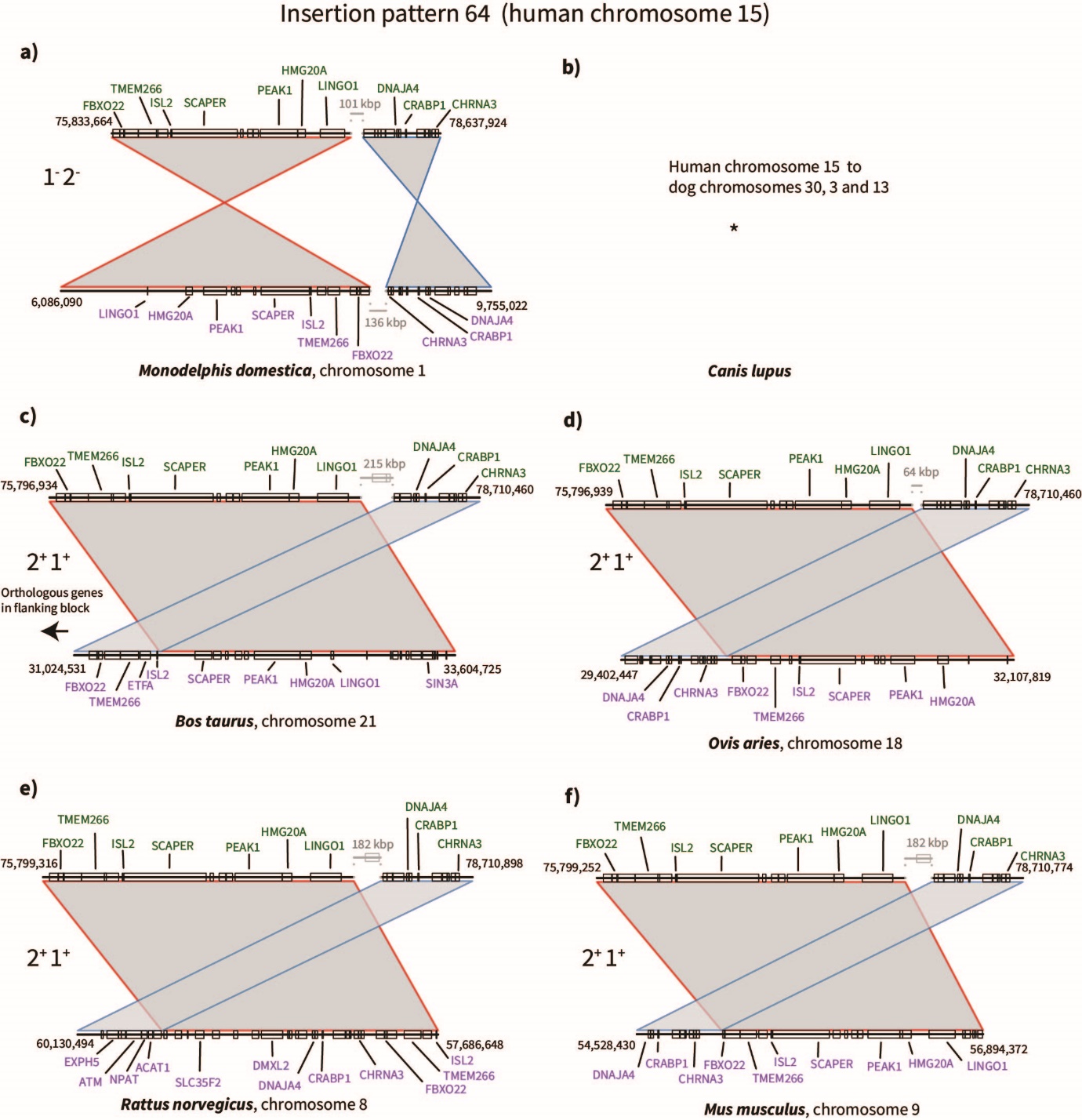

**Figure S2. Insertion pattern 64. No synteny map is shown in panel b) because the pattern was degenerated and found in three separate chromosomes in dog.**

##### Insertion patterns 67 for primates

The pattern was found in opossum, dog, cow, sheep, rat and mouse (Figure S3). The opossum synteny blocks were somewhat degenerated and had a predicted third block (in blue) in between the two blocks (red and green) compared to the remaining species. The predicted third block for opossum contained genes that were in gaps (outside of the synteny blocks) for the remaining organisms. For rat, none of the gene annotations were clear orthologs to the human genes in the area.

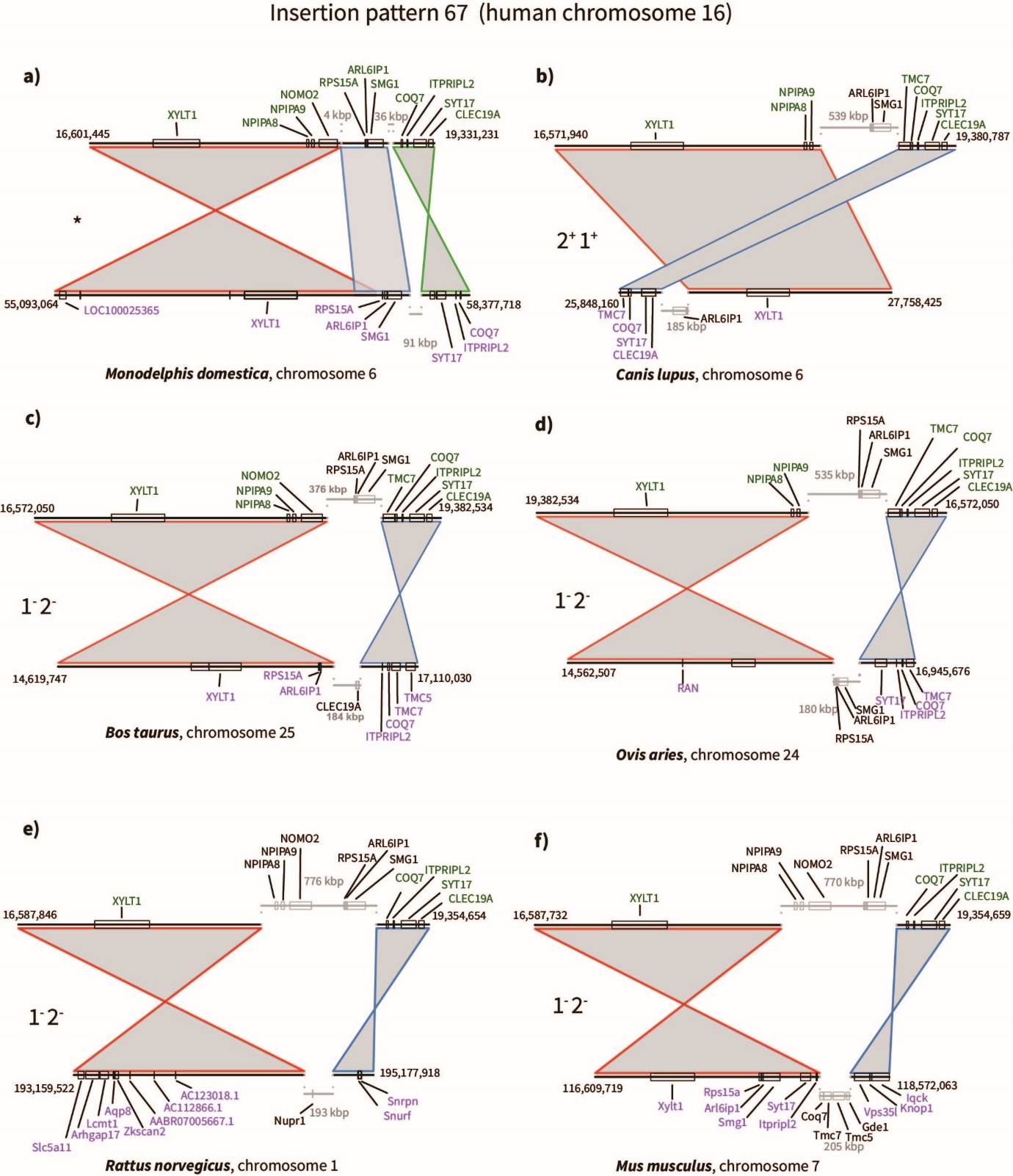

**Figure S3. Insertion pattern 67.**

##### Insertion patterns 66 for human and chimpanzee

The pattern was found for opossum, dog, cow, sheep, rat, mouse and rhesus monkey, but was somewhat degraded for cow and rat (Figure S4). For cow, the pattern appeared to be degraded/rearranged within the same chromosome. Three blocks were found covering the same human area as for the other insertion patterns. In cow the area was split up in four blocks, with one additional block falling in between two (block 2 and 4). In addition, a flanking block further upstream in cow, block 0, was identified to contain all the same gene annotations as for human in block 4. Block 0 was predicted to have homology to human downstream in the same chromosome.

The coordinates for all blocks are given below:

Block 1 – Human chr15: 81,480,549- 82,287,834; Cow chr21:23,955,027- 24544033(+)

Block 2 - Human chr15: 82,536,244- 83015387; Cow chr21:23,300,928- 23,729,416 (+)

Block 3 - Human chr15: 83,016,018- 84,150,500; Cow chr21: 24,458,072-25,519,818(-)

Block 4 - Human chr15: 84,598,057- 85,169,524; Cow chr21: 22,698,129-23,297,618 (+)].

Block 0 – Human chr15: 90,314,298- 91,022,692; Cow chr21: 22,071,827- 22,698,143 (-)

Block 1-4 were plotted to show the pattern. The arrow in Figure S8 indicates the block with gene annotations matching human block 4.

For rat, none of the human genes annotated in block 1 had orthologs in the area predicted. All orthologs were found flanking downstream of the predicted homology. The genes *ZPF592*, *ALPK3*, *SLC28A1* where found in the downstream block, whereas *PDE8A* was immediate upstream and not allocated to the flanking block in the synteny prediction.

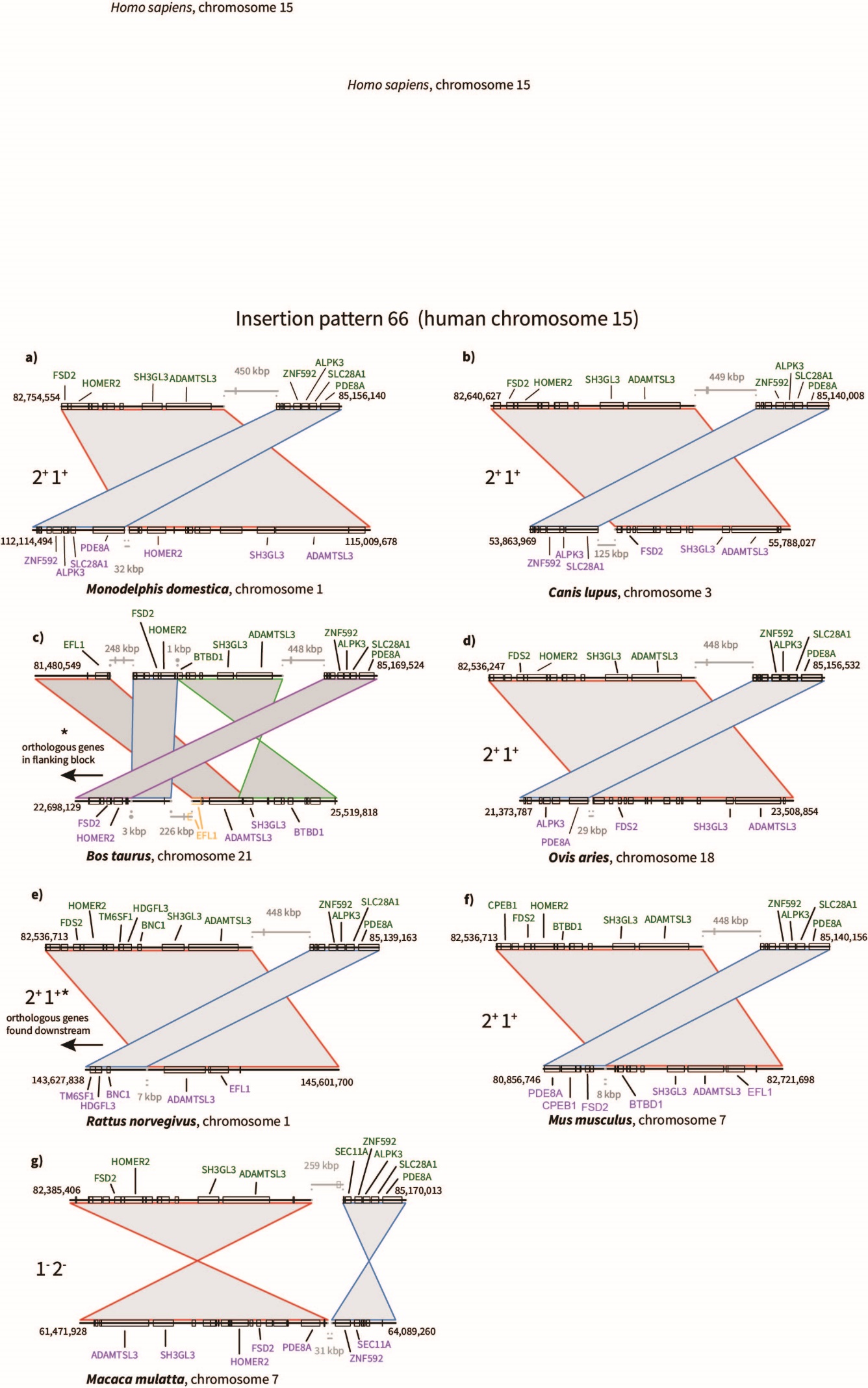

**Figure S4. Insertion pattern 66.**

##### Insertion pattern 75 for human and chimpanzee

The pattern was found for opossum, dog, cow, sheep, rat and mouse, but was degraded/rearranged for rat (Figure S5). For rat, the orthologous genes *UTP6* and *SUZ12* were found at the intercept between two blocks and downstream of the predicted synteny block pairs. This suggests that the synteny software incorrectly called the blocks

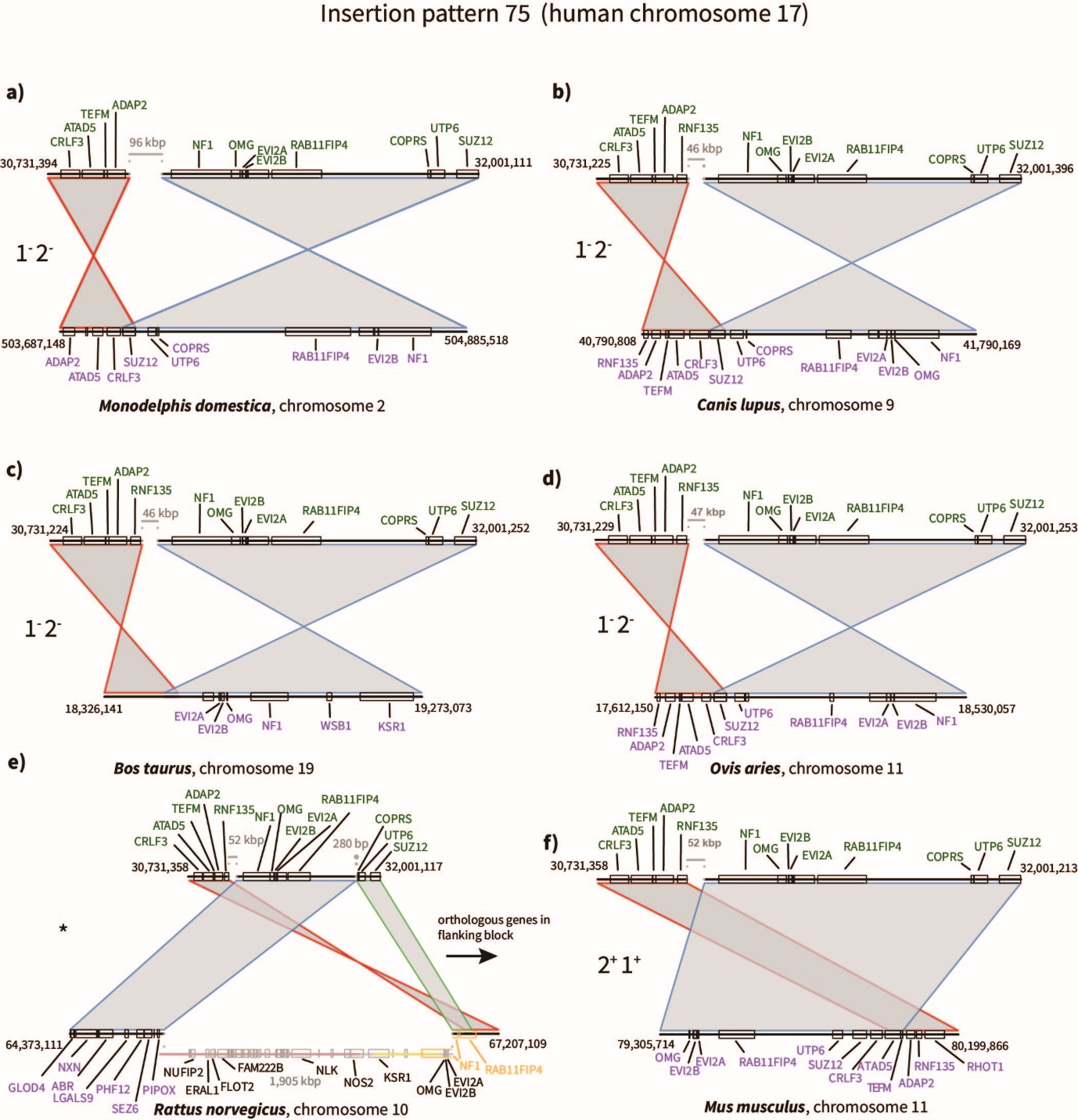

**Figure S5. Insertion pattern 75.**

#### Large canonical circle insertion patterns in ungulates

The largest and most frequent circle insertion shared between species were found for the ungulates cow and sheep. Below we draw examples of three large circle insertion patterns (22, 56 and 77; 13-43 Mb) between the ungulates cow and sheep (Figure S6-S8).

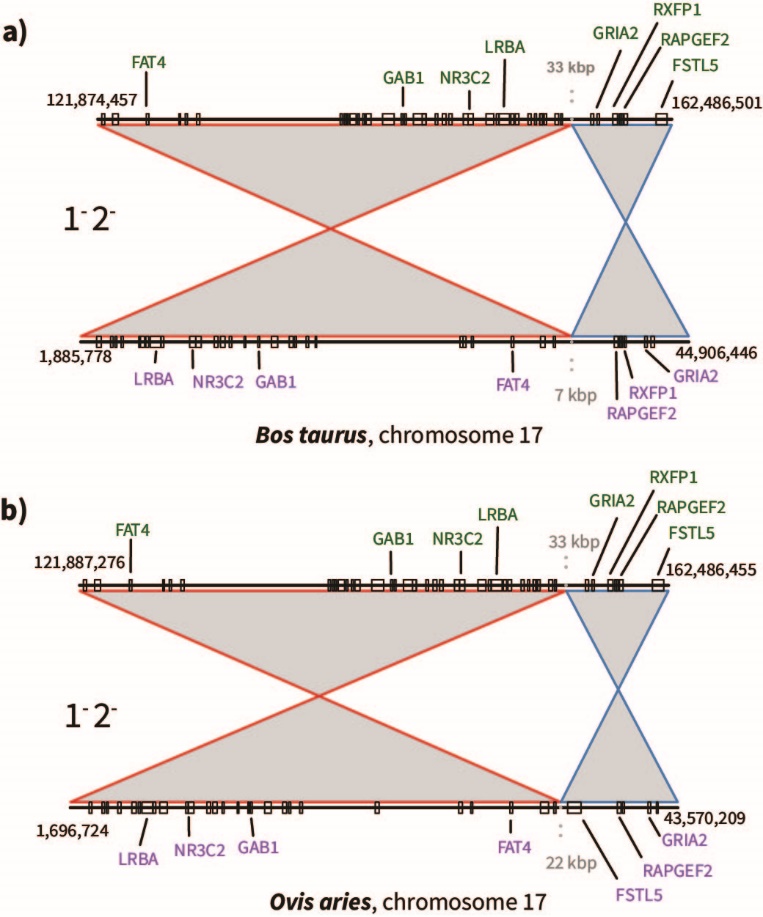

**Figure S6. Circle insertion pattern 22 between cow and sheep.**

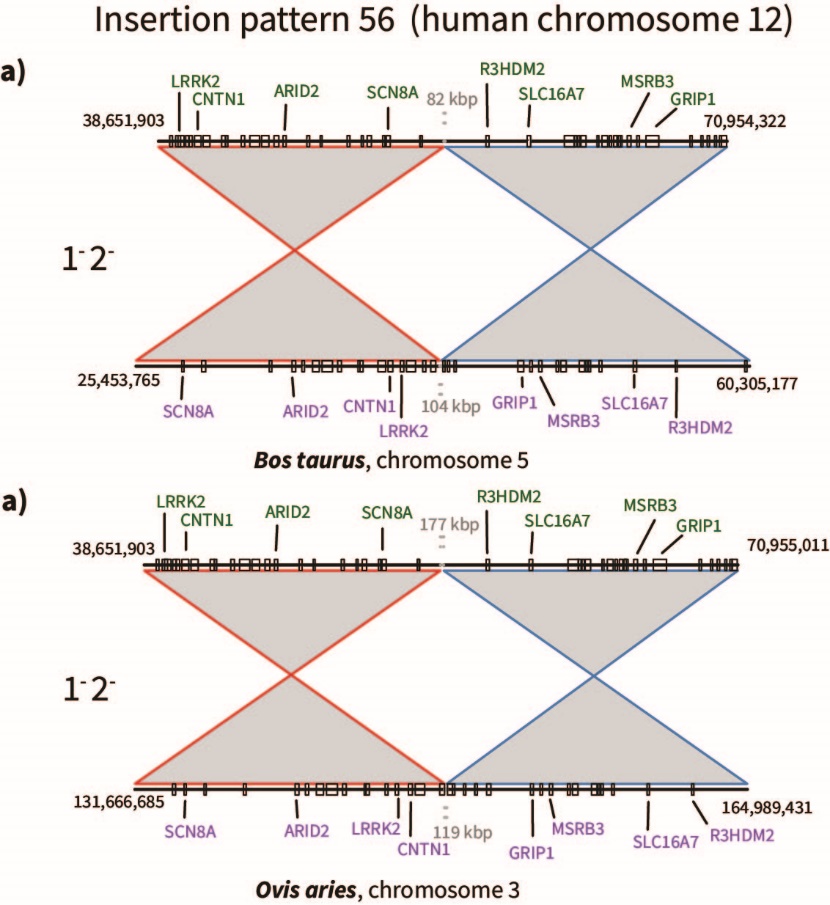

**Figure S7. Circle insertion pattern 56**

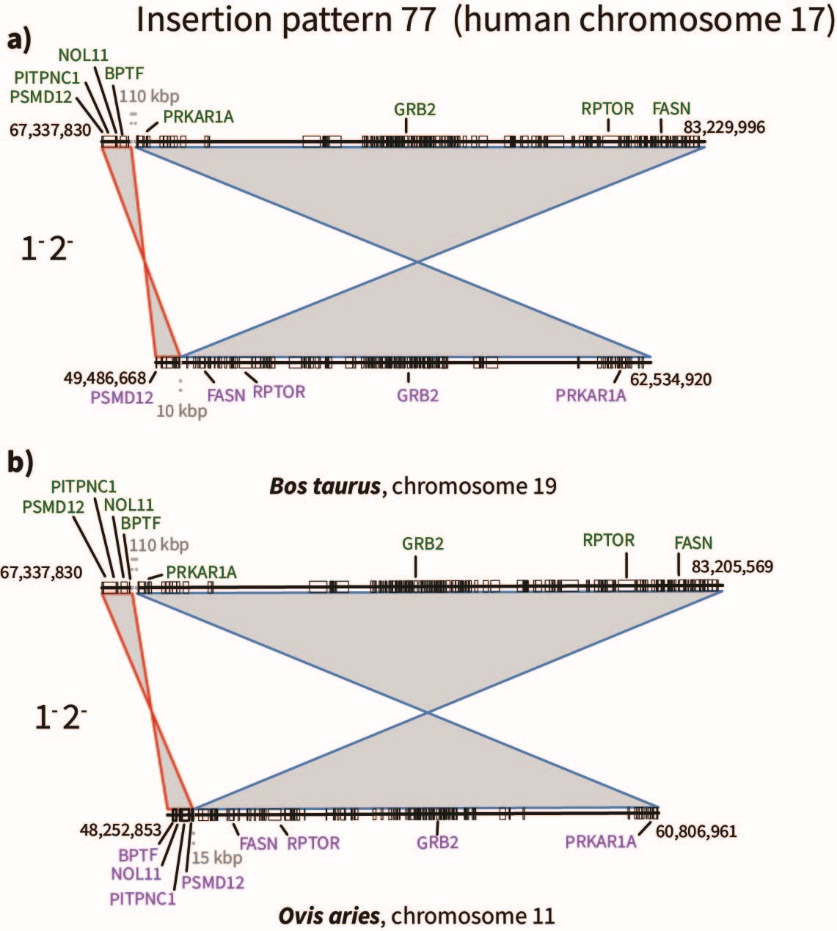

**Figure S8. Circle insertion pattern 77.**

#### Circle insertion pattern for dog, cow and sheep

One insertion pattern (74) was observed at the branch between cow, sheep and dog. For **cow**, all genes in block 1 (*TTC19*, *NCOR1*, *PIGL*, *TRPV2*) were found upstream, suggesting a degenerated pattern.

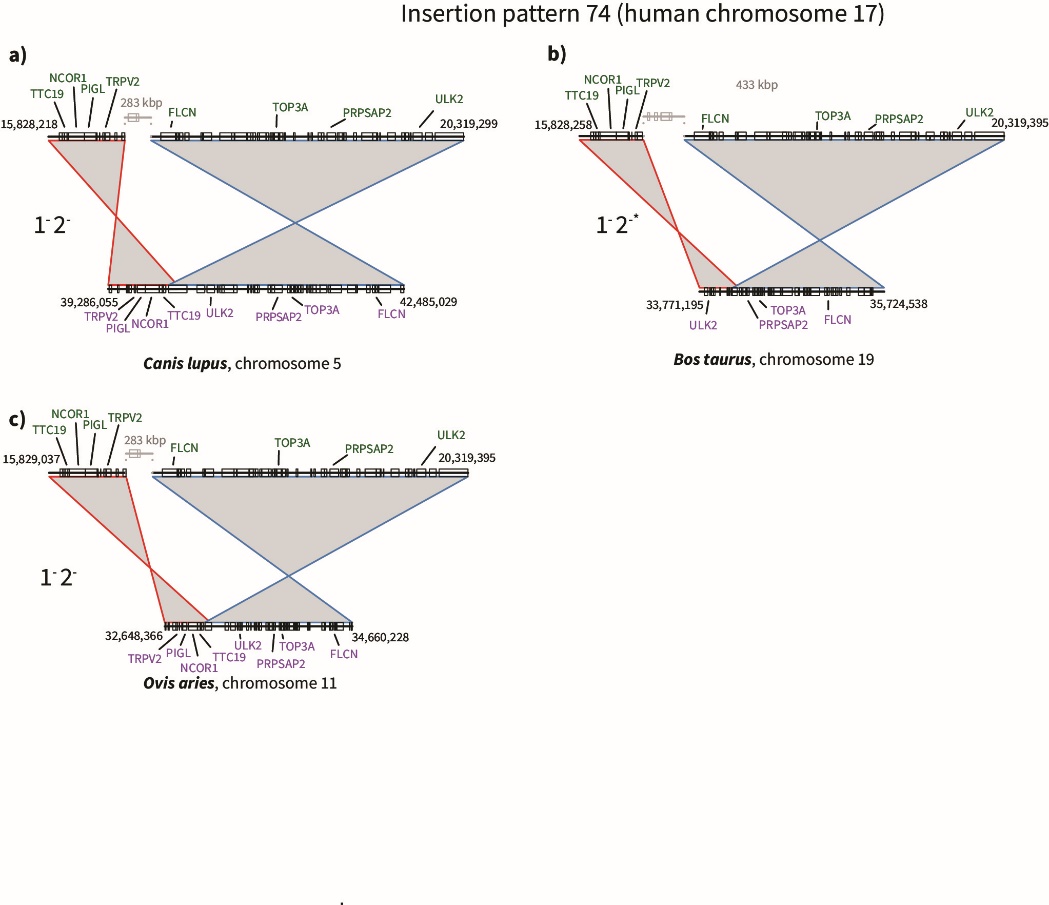

**Figure S9. Circle insertion pattern 74**

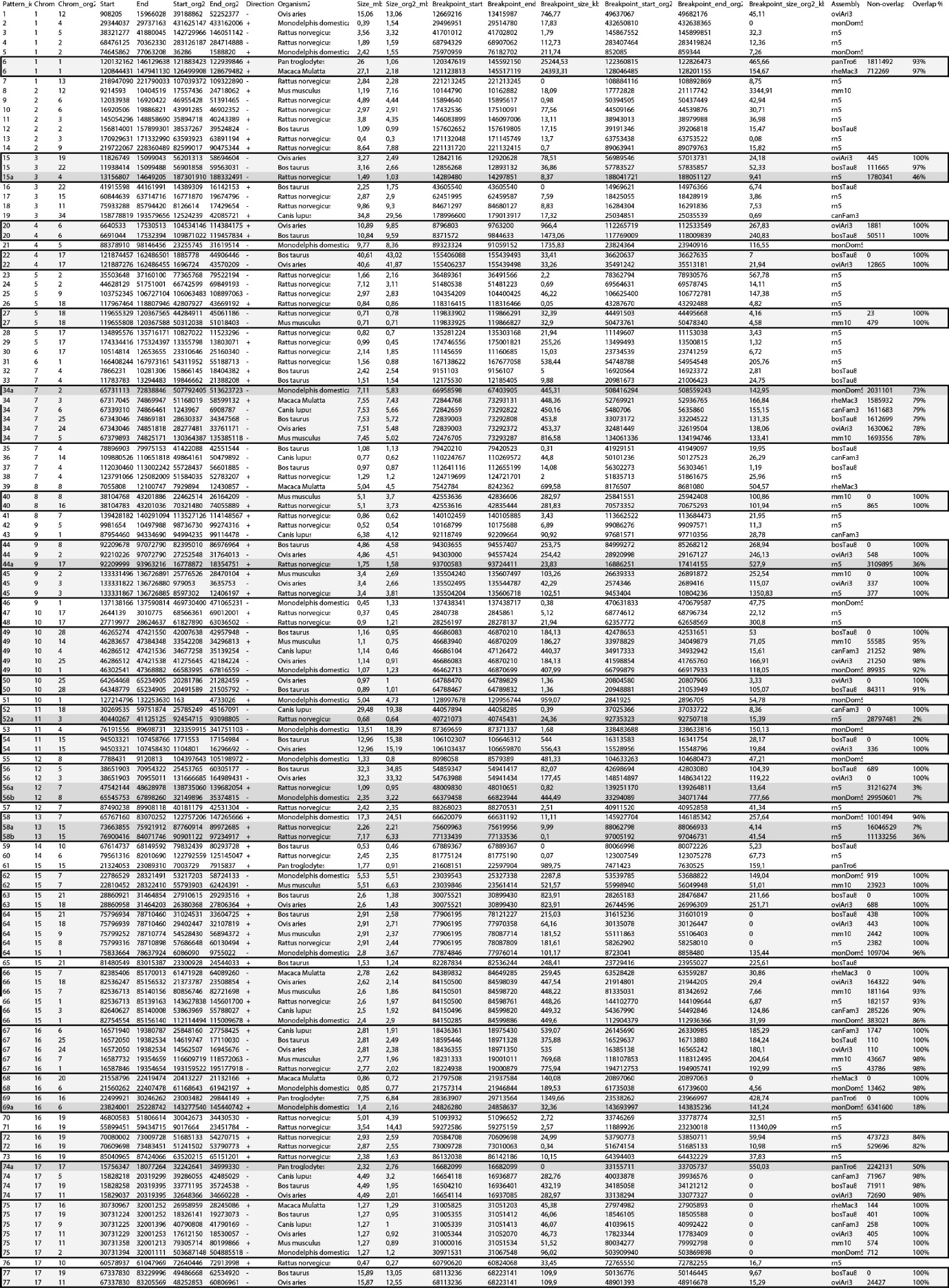

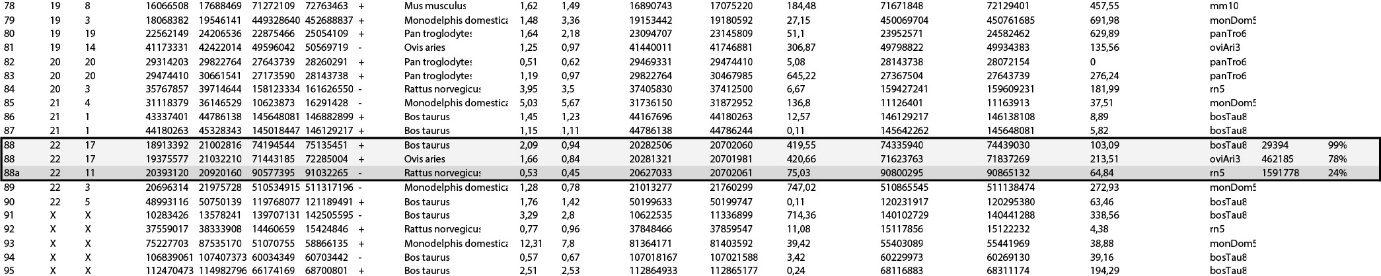

**Table S1. Properties for all circle insertion patterns. Patterns that are overlapping, are indicated with boxes**

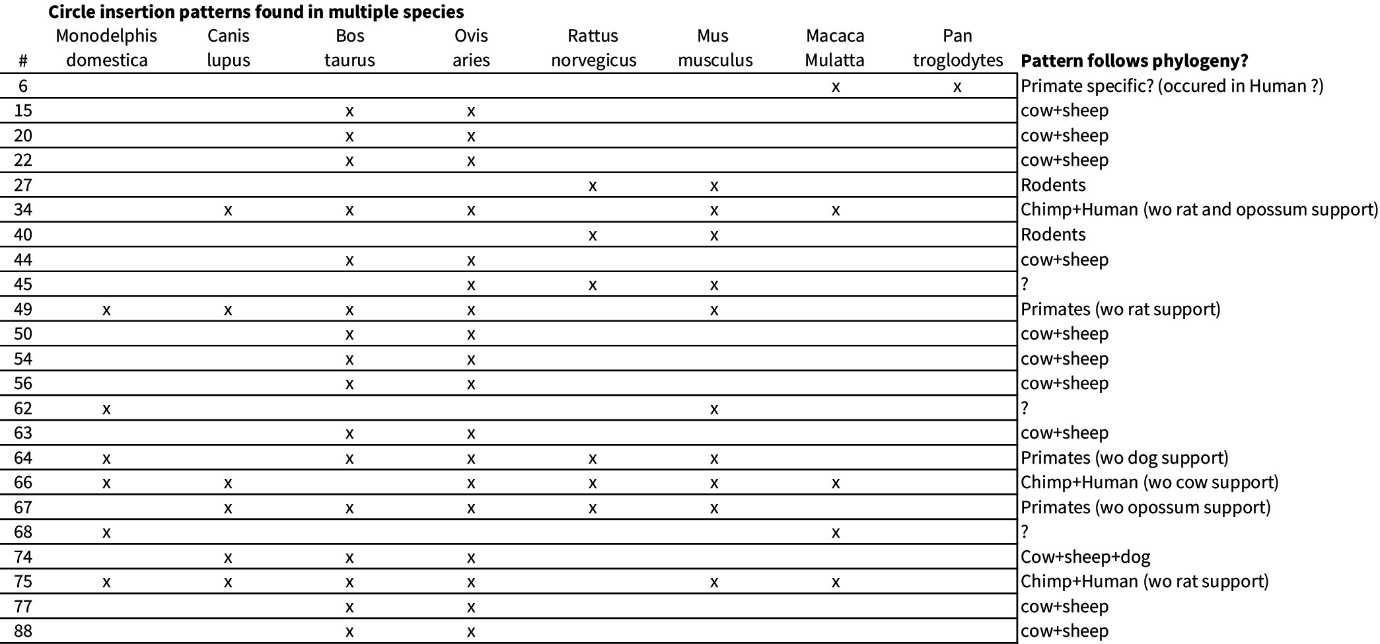

**Table S2. Comparison of circle insertion patterns found in multiple species**

**Occurrence of random circle insertion patterns**

In order to test whether the patterns were occurring by complete randomness, we randomized the synteny maps 10,000 times for the eight mammal species and extracted randomly generated circle insertion patterns (Figure S10). For all species, the number of circle insertion patterns is higher than the randomized number.

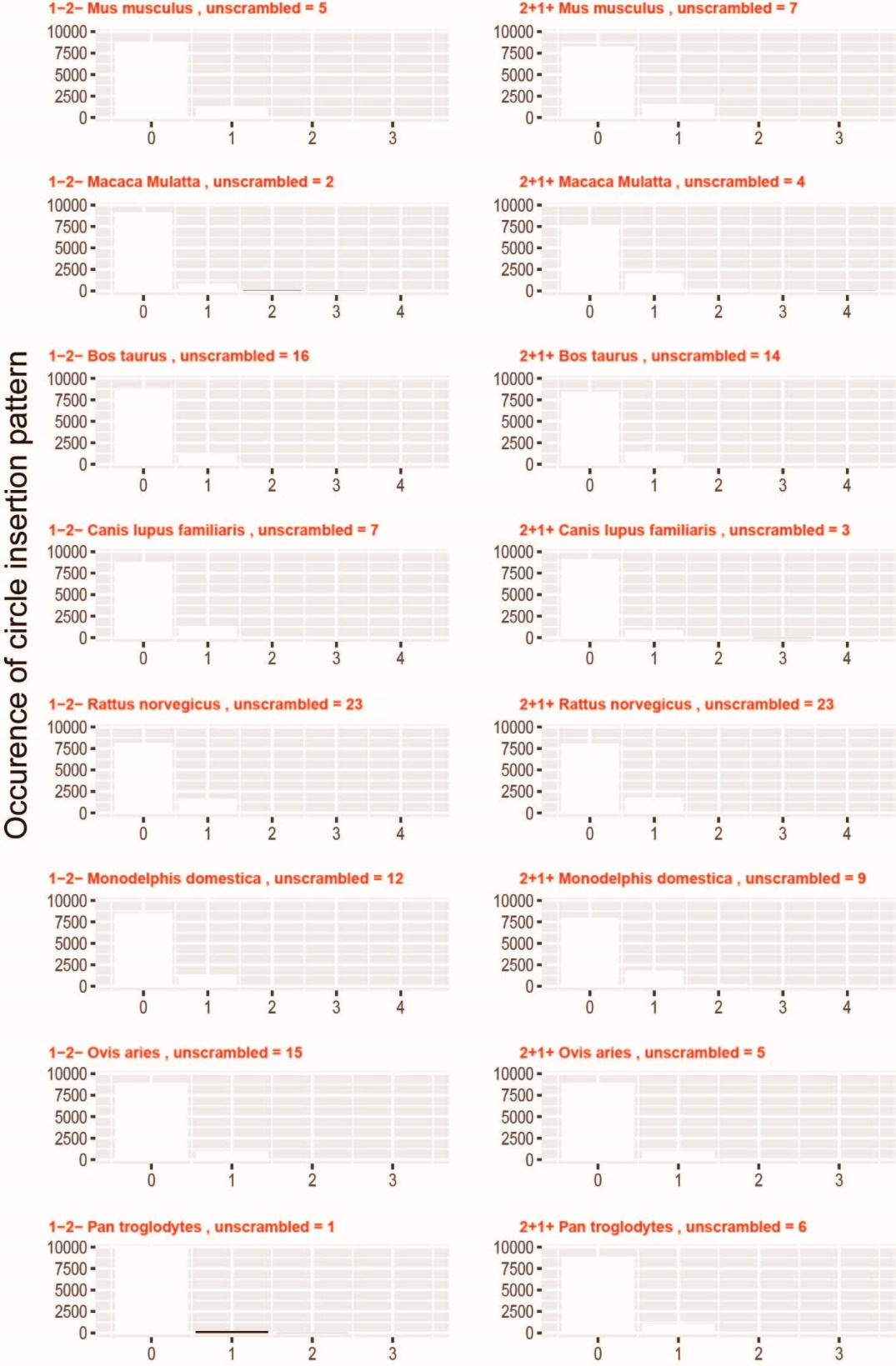

**Figure S10: Circle insertion patterns found in 10,000 randomized synteny blocks.** Numbers in red, labelled “unscrambled” indicate the number of circle insertion patterns found in the real synteny maps.
